# N4-hydroxycytidine and inhibitors of dihydroorotate dehydrogenase synergistically suppress SARS-CoV-2 replication

**DOI:** 10.1101/2021.06.28.450163

**Authors:** Kim M. Stegmann, Antje Dickmanns, Natalie Heinen, Uwe Groß, Dirk Görlich, Stephanie Pfaender, Matthias Dobbelstein

## Abstract

Effective therapeutics to inhibit the replication of SARS-CoV-2 in infected individuals are still under development. The nucleoside analogue N4-hydroxycytidine (NHC), also known as EIDD-1931, interferes with SARS-CoV-2 replication in cell culture. It is the active metabolite of the prodrug Molnupiravir (MK-4482), which is currently being evaluated for the treatment of COVID-19 in advanced clinical studies. Meanwhile, inhibitors of dihydroorotate dehydrogenase (DHODH), by reducing the cellular synthesis of pyrimidines, counteract virus replication and are also being clinically evaluated for COVID-19 therapy. Here we show that the combination of NHC and DHODH inhibitors such as teriflunomide, IMU-838/vidofludimus, and BAY2402234, strongly synergizes to inhibit SARS-CoV-2 replication. While single drug treatment only mildly impaired virus replication, combination treatments reduced virus yields by at least two orders of magnitude. We determined this by RT-PCR, TCID_50_, immunoblot and immunofluorescence assays in Vero E6 and Calu-3 cells infected with wildtype and the Alpha and Beta variants of SARS-CoV-2. We propose that the lack of available pyrimidine nucleotides upon DHODH inhibition increases the incorporation of NHC in nascent viral RNA, thus precluding the correct synthesis of the viral genome in subsequent rounds of replication, thereby inhibiting the production of replication competent virus particles. This concept was further supported by the rescue of replicating virus after addition of pyrimidine nucleosides to the media. Based on our results, we suggest combining these drug candidates, which are currently both tested in clinical studies, to counteract the replication of SARS-CoV-2, the progression of COVID-19, and the transmission of the disease within the population.

**SIGNIFICANCE:**

- The strong synergy displayed by DHODH inhibitors and the active compound of Molnupiravir might enable lower concentrations of each drug to antagonize virus replication, with less toxicity.
- Both Molnupiravir and DHODH inhibitors are currently being tested in advanced clinical trials or are FDA-approved for different purposes, raising the perspective of rapidly testing their combinatory efficacy in clinical studies.
- Molnupiravir is currently a promising candidate for treating early stages of COVID-19, under phase II/III clinical evaluation. However, like Remdesivir, it appears only moderately useful in treating severe COVID-19. Since the combination inhibits virus replication far more strongly, and since DHODH inhibitors may also suppress excessive immune responses, the combined clinical application bears the potential of alleviating the disease burden even at later stages.

## INTRODUCTION

In order to combat the COVID-19 pandemic, vaccines have successfully been established, while efficient therapeutics are still under development (Doherty, 2021). At present, the suppression of excessive inflammatory and immune responses by steroids has been successfully applied in the clinics (Horby et al., 2021; Tomazini et al., 2020). However, this strategy does not directly interfere with virus replication. So far, the only FDA-approved antiviral drug for the treatment of COVID-19, the adenosine analogue Remdesivir, has only been moderately successful in shortening hospitalization by a few days, with variations reported between clinical studies (Beigel et al., 2020; Goldman et al., 2020). Antiviral nucleoside analogues typically antagonize the synthesis of viral genomes. Upon entry into the host cell, they are triphosphorylated at their 5’ positions. Subsequently, they interfere with the activity of the viral nucleic acid polymerase or compromise the function of the newly synthesized viral genomes (Pruijssers and Denison, 2019).

Next to Remdesivir, another nucleoside analogue showed promising performance against SARS-CoV-2. Molnupiravir, also known as EIDD-2801 or MK-4482, is the prodrug of N4-hydroxycytidine (NHC), or EIDD-1931 (Cox et al., 2021; Painter et al., 2021; Sheahan et al., 2020; Wahl et al., 2020; Wahl et al., 2021). NHC differs from cytidine by a hydroxyl group at the pyrimidine base. According to current literature, this does not impair the incorporation of triphosphorylated NHC into nascent RNA by the viral RNA polymerase. However, due to a tautomeric interconversion within the NHC base, RNA strands with incorporated NHC lead to erroneous RNA replication when used as a template (Jena, 2020). NHC can basepair with guanosine, but also with adenosine, thus leading to multiple errors in the subsequently synthesized viral RNA genomes resulting in replication-deficient virus particles. Molnupiravir has been found to be active against SARS-CoV-2 replication in in vitro and in vivo (Sheahan et al., 2020; Wahl et al., 2021). It has been shown to also prevent SARS-CoV-2 transmission in vivo (Cox et al., 2021), and its clinical efficacy is currently being investigated in large-scale phase II clinical trials (Painter et al., 2021), NCT04575584, NCT04575597, NCT04405739.

Besides immunosuppression and direct interference with virus replication, an alternative approach of treatment against SARS-CoV-2 aims at reducing the cellular synthesis of nucleotides, thereby indirectly impairing the synthesis of viral RNA. We (Stegmann et al., 2021) and others (Caruso et al., 2021; Zhang et al., 2021) have previously reported the high demand on cellular nucleotide biosynthesis during SARS-CoV-2 infection, resulting in an antiviral efficacy of folate antagonists that impair purine synthesis. Moreover, in the context of nucleotide biosynthesis, the inhibition of dihydroorotate dehydrogenase (DHODH) represents an attractive strategy to antagonize SARS-CoV-2 replication. DHODH catalyzes a key step during the pyrimidine synthesis. Unlike all other cytosolic enzymes involved in this metabolic pathway, DHODH localizes to the inner mitochondrial membrane, where it transfers reduction equivalents from dihydroorotate to ubiquinone moieties of the respiration chain. As a result, orotate becomes available for the subsequent synthesis steps to obtain uridine monophosphate and later cytidine triphosphate. A number of DHODH inhibitors have become available for clinical testing or were even approved for therapy (Munier-Lehmann et al., 2013). Mostly, DHODH inhibitors are used for immunosuppression, presumably because of their ability to reduce the proliferation of B and T cells. Recently, however, some DHODH inhibitors were successfully tested with regard to their efficacy in preventing the replication of viruses (Hoffmann et al., 2011; Zhang et al., 2012), including SARS-CoV-2 (Hahn et al., 2020; Luban et al., 2021; Xiong et al., 2020). One DHODH inhibitor, IMU-838 (vidofludimus), was further clinically evaluated for COVID-19 therapy and was found effective according to secondary criteria e.g. time to clinical improvement or viral burden (CALVID-1, trial identifier NCT04379271).

We therefore hypothesized that the suppression of pyrimidine synthesis should increase the ratio of NHC triphosphate vs. cytidine triphosphate in the infected cell, thus enhancing the incorporation of NHC into the viral RNA, and resulting in the production of replication deficient viral particles. We found that the combination of NHC and DHODH inhibitors resulted in profoundly synergistic suppression of SARS-CoV-2 replication, presenting a potential treatment strategy that could be exploited in the clinic.

## MATERIALS & METHODS

### Experimental Model And Methods

#### Cell culture

Vero E6 cells (Vero C1008) were maintained in Dulbecco’s modified Eagle’s medium (DMEM with GlutaMAX™, Gibco) supplemented with 10% fetal bovine serum (Merck), 50 units/ml penicillin, 50 μg/ml streptomycin (Gibco), 2 µg/ml tetracycline (Sigma) and 10 µg/ml ciprofloxacin (Bayer) at 37°C in a humidified atmosphere with 5% CO_2_. Calu-3 cells were maintained in Eagle’s Minimum Essential Medium (EMEM, ATCC) supplemented with 10% fetal bovine serum and penicillin/streptomycin. Cells were authenticated by the Leibniz-Institute DSMZ and routinely tested and ensured to be negative for mycoplasma contamination.

#### Treatments and SARS-CoV-2 infection

30,000 cells per well were seeded into 24-well-plates using medium containing 2% fetal bovine serum (FBS) and incubated for 8 hours at 37 °C. Cells were treated with β-D-N^4^-Hydroxycytidine (NHC/EIDD-1931, Cayman Chemical 9002958), IMU-838 (Immunic Therapeutics), BAY2402234 (Selleckchem S8847), Teriflunomide (Selleckchem S4169), ASLAN003 (Selleckchem S9721), Brequinar (Selleckchem S6626), Uridine (Selleckchem S2029) and Cytidine (Selleckchem S2053) at the concentrations indicated in the figure legends. When preparing stock solutions, all compounds were dissolved in DMSO. After 24 hours, the cells were infected with virus stocks corresponding to 1*10^7^ RNA-copies of SARS-CoV-2 (= 30 FFU) and incubated for 48 hours at 37 °C, as described (Stegmann et al., 2021). Images of mock and SARS-CoV-2 infected cells were taken by brightfield microscopy. SARS-CoV-2 ‘wildtype’ was isolated from a patient sample taken in March 2020 in Göttingen, Germany (Stegmann et al., 2021). The SARS CoV-2 variants Alpha and Beta were kindly provided by the Robert Koch Institute at Berlin, Germany.

To determine the Median Tissue Culture Infectious Dose (TCID_50_) per mL, 30,000 Vero E6 cells per well were pre-treated with NHC, IMU-838 or the indicated combinations for 24 hours, infected with SARS-CoV-2 strains Wuhan, B.1.1.7/Alpha or B.1.351/Beta (MOI 0.1) for 1 hour and again treated with the drugs for additional 48 hours. The virus-containing supernatant was titrated (end point dilution assay) to calculate the TCID_50_/mL.

#### Quantitative RT-PCR for virus quantification

For RNA isolation, the SARS-CoV-2 containing cell culture supernatant was mixed (1:1 ratio) with the Lysis Binding Buffer from the Magnapure LC Kit # 03038505001 (Roche) to inactivate the virus. The viral RNA was isolated using Trizol LS, chloroform, and isopropanol. After washing the RNA pellet with ethanol, the isolated RNA was resuspended in nuclease-free water. Quantitative RT-PCR was performed according to a previously established RT-PCR assay involving a TaqMan probe (Corman et al., 2020), to quantify virus yield. The following oligonucleotides were used for qRT-PCR, which amplify a genomic region corresponding to the envelope protein gene (26,141 – 26,253), as described (Corman et al., 2020).

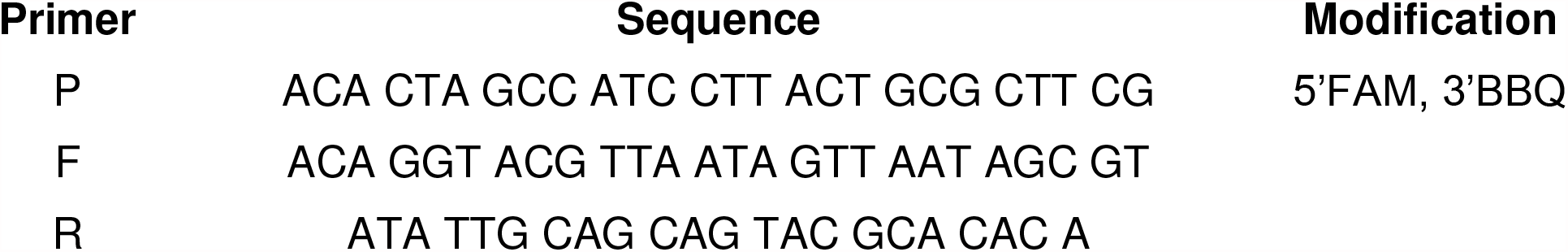

The amount of SARS-CoV-2 RNA determined upon infection without any treatment was defined as 100%, and the other RNA quantities were normalized accordingly.

#### Immunofluorescence analyses

Vero E6 cells were seeded onto 8-well chamber slides (Nunc) and treated/infected as indicated. After 48 hours of SARS-CoV-2 infection, the cells were washed once in PBS and fixed with 4% formaldehyde in PBS for 1 hour at room temperature. After permeabilization with 0.5% Triton X-100 in PBS for 30 minutes and blocking in 10% FBS/PBS for 10 minutes, primary antibodies were used to stain the SARS-CoV-2 Nucleoprotein (N; Sino Biological #40143-R019, 1:8000) and Spike protein (S; GeneTex#GTX 632604, 1:2000) overnight. The secondary Alexa Fluor 546 donkey anti-rabbit IgG and Alexa Fluor 488 donkey anti-mouse IgG (Invitrogen, 1:500, diluted in blocking solution) antibodies were added together with 4′,6-diamidino-2-phenylindole (DAPI) for 1.5 hours at room temperature. Slides with cells were mounted with Fluorescence Mounting Medium (DAKO) and fluorescence signals were detected by microscopy (Zeiss Axio Scope.A1).

#### Immunoblot analysis

Cells were washed once in PBS and harvested in radioimmunoprecipitation assay (RIPA) lysis buffer (20 mM TRIS-HCl pH 7.5, 150 mM NaCl, 10 mM EDTA, 1% Triton-X 100, 1% deoxycholate salt, 0.1% SDS, 2 M urea), supplemented with protease inhibitors. Samples were briefly sonicated and protein extracts quantified using the Pierce BCA Protein assay kit (Thermo Fisher Scientific). After equalizing the amounts of protein, samples were incubated at 95 °C in Laemmli buffer for 5 min and separated by sodium dodecyl sulfate polyacrylamide gel electrophoresis (SDS-PAGE). To determine the presence of viral proteins, the separated proteins were transferred to a nitrocellulose membrane, blocked in 5% (w/v) non-fat milk in TBS containing 0.1% Tween-20 for 1 hour, and incubated with primary antibodies at 4 °C overnight, followed by incubation with peroxidase-conjugated secondary antibodies (donkey anti-rabbit or donkey anti-mouse IgG, Jackson Immunoresearch). The SARS-CoV-2 Spike- and Nucleoprotein, DHODH and HSC70 (loading control) were detected using either Super Signal West Femto Maximum Sensitivity Substrate (Thermo Fisher) or Immobilon Western Substrate (Millipore).

| **Antibody** | **Source (Catalogue number)** | **Concentration** |
| --- | --- | --- |
| SARS-CoV-2 Spike | GeneTex GTX 632604 | 1:1,000 |
| SARS-CoV-2 Nucleoprotein | Sino Biological 40143-R019 | 1:5,000 |
| DHODH | Santa Cruz sc-166348 | 1:250 |
| HSC70 | Santa Cruz sc-7298 | 1:10,000 |

#### Quantification of LDH release to determine cytotoxicity

3,500 Vero E6 cells were seeded into 96-well-plates and treated with DHODH inhibitors and/or NHC as indicated, for 72 hours (corresponding to the incubation of cells with drugs before and after infection). The release of lactate dehydrogenase (LDH) into the cell culture medium was quantified by bioluminescence using the LDH-Glo™ Cytotoxicity Assay kit (Promega). 10% Triton X-100 was added to untreated cells for 15 minutes to determine the maximum LDH release, whereas the medium background (= no-cell control) served as a negative control. Percent cytotoxicity reflects the proportion of LDH released to the media compared to the overall amount of LDH in the cells, and was calculated using the following formula:

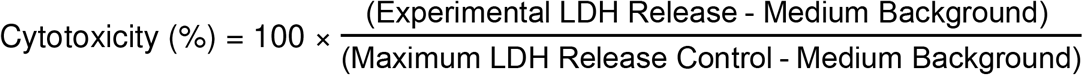

#### Quantification and statistical analysis

Statistical testing was performed using Graph Pad Prism 6 (RRID:SCR_002798). A two-sided unpaired Student’s t-test was calculated, and significance was assumed where p-values ≤ 0.05. Asterisks represent significance in the following way: ^****^, p ≤ 0.0001; ^***^, p ≤ 0.005; ^**^, p ≤ 0.01; ^*^, p ≤ 0.05.

## RESULTS

### NHC and DHODH inhibitors synergistically interfere with SARS-CoV-2 replication, without signs of cytotoxicity

The biosynthesis of pyrimidines is crucial for RNA replication (Fig. 1A). The enzyme dihydroorotate dehydrogenase (DHODH) catalyzes the oxidation of dihydroorotate to orotate, which is a precursor of cytidine triphosphate (CTP). In the presence of N4-hydroxycytidine (NHC), its active metabolite NHCTP competes with CTP for incorporation into nascent RNA. We hypothesized that the suppression of cellular CTP synthesis by DHODH inhibitors favors the incorporation of NHCTP into newly synthesized SARS-CoV-2 RNA, and thus potentiates the antiviral efficacy of NHC. To test this, we combined both drugs for treatment of Vero E6 or Calu-3 cells, followed by infection with SARS-CoV-2. We applied suboptimal concentrations of NHC and the DHODH inhibitors BAY2402234, teriflunomide and IMU-838 to moderately diminish virus replication. We treated cells with the drugs at these concentrations, alone or in combination, 24 hours before and during infection with SARS-CoV-2. At these concentrations, neither NHC nor DHODH inhibitors grossly affected the development of a cytopathic effect (CPE) caused by SARS-CoV-2. Strikingly, however, the combination of NHC and DHODH inhibitors was far more efficient in preventing the CPE (Fig. 1B) and reducing virus yield, as determined by the Median Tissue Culture Infectious Dose (TCID_50_/mL) (Fig. 1C). Combining the drugs did not produce morphologic signs of cytotoxicity in non-infected cells (Fig. 1B). To quantify a possible increase in cytotoxicity, we measured the release of lactate dehydrogenase (LDH) into the culture supernatant. Compared to the DMSO control, neither the single treatments nor the combinations displayed major cytotoxicity (Fig. 1D). Hence, the drug combination synergistically interferes with CPE and virus yield, without displaying cytotoxicity on its own.

**FIGURE 1:**
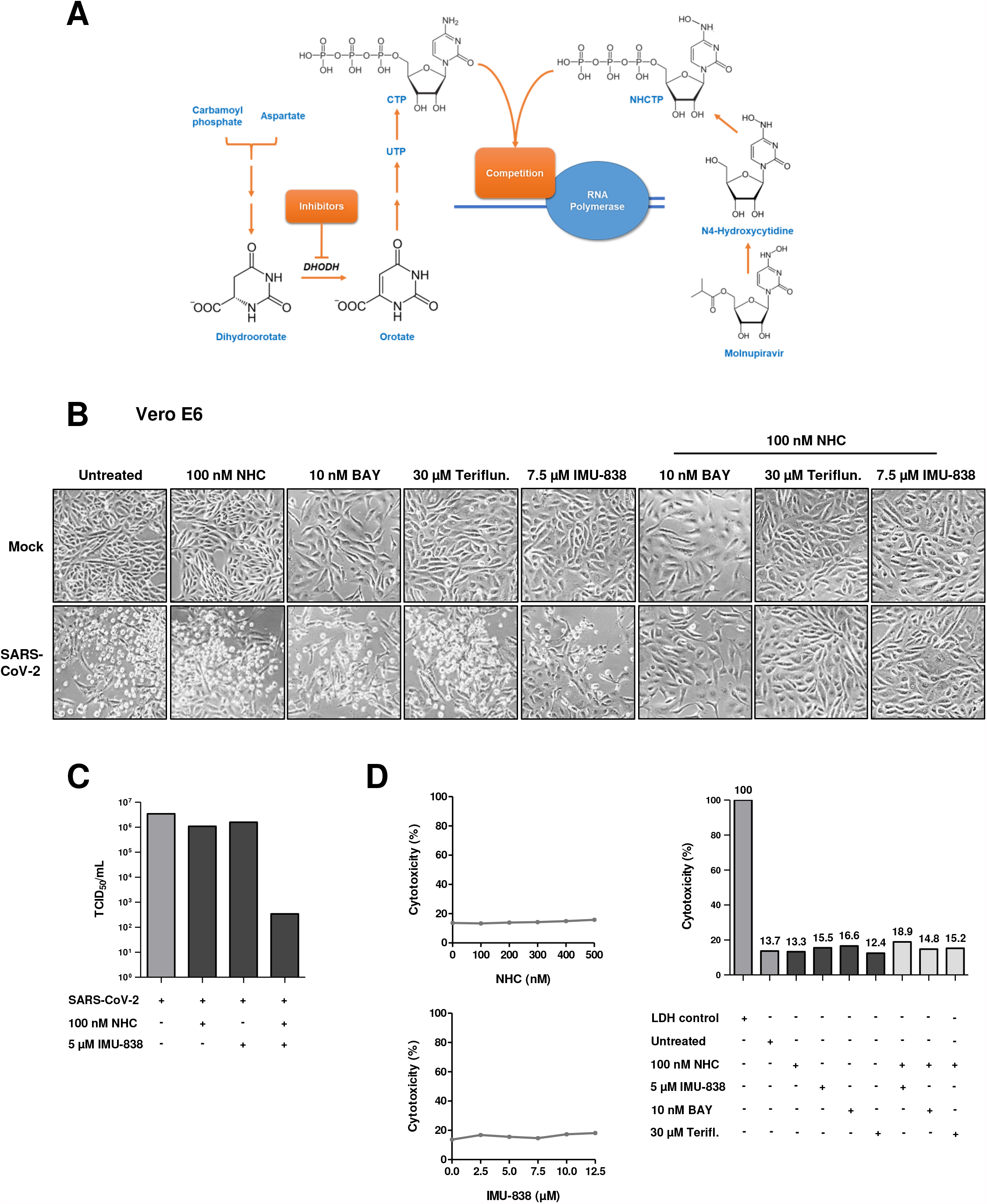
The combination of N4-hydroxycytidine (NHC) and inhibitors of dihydroorotate dehydrogenase (DHODH) strongly impairs SARS-CoV-2 replication without detectable cytotoxicity. **(A)** Mechanistic concept for the synergistic effect of DHODH inhibitors and N4-hydroxycytidine (NHC) in counteracting SARS-CoV-2 RNA replication. Biosynthesis of pyrimidines starts with carbamoyl phosphate and aspartate to form dihydroorotate in a series of reactions. Dihydroorotate is further oxidized to orotate by dihydroorotate dehydrogenase (DHODH) and later converted to uridine triphosphate (UTP) and cytidine triphosphate (CTP). Molnupiravir is the prodrug of NHC, which is further converted to the corresponding triphosphate (NHCTP), which competes with CTP for incorporation into nascent virus RNA. The suppression of CTP synthesis by inhibitors of DHODH is expected to enhance the incorporation of NHCTP into the viral RNA and the false incorporation of nucleotides in subsequent rounds of replication. **(B)** Reduced cytopathic effect (CPE) by NHC and DHODH inhibitors. Vero E6 cells were treated with drugs or the DMSO control for 24 hrs, inoculated with SARS-CoV-2, and further incubated in the presence of the same drugs for 48 hrs. Cell morphology was assessed by brightfield microscopy. Note that the CPE was readily visible when comparing mock-infected and virus-infected cells, but only to a far lesser extent when the cells had been incubated with both NHC and DHODH inhibitors. **(C)** Reduction of the Median Tissue Culture Infectious Dose (TCID_50_) by NHC and DHODH inhibitors. Vero E6 cells were treated with NHC, IMU-838 or the combination of both compounds for 24 hrs before infection, and then throughout the time of infection. Cells were infected with SARS-CoV-2 wildtype (MOI 0.1) and further incubated for 48 hrs. The supernatant was titrated to determine the TCID_50_/mL. **(D)** Lack of measurable cytotoxicity by NHC and DHODH inhibitors. Vero E6 cells were treated with NHC and/or IMU-838, BAY2402234 and teriflunomide at the indicated concentrations for 72 hrs. The release of lactate dehydrogenase (LDH) to the supernatant was quantified by bioluminescence as a read-out for cytotoxicity. The percentages reflect the proportion of LDH released to the media, compared to the overall amount of LDH in the cells (LDH control).

### Combinations of NHC and DHODH inhibitors distinctly reduce viral RNA yield upon infection with SARS-CoV-2

We combined different concentrations of the DHODH inhibitor IMU-838 and NHC, and determined their impact on the release of viral RNA, along with the combination index (Chou, 2010). When combined, the effective concentrations of IMU-838 and NHC were between 5-10 µM and 100-300 nM, respectively (Fig. 2A). Moreover, we combined the DHODH inhibitors BAY2402234, teriflunomide, ASLAN003 and brequinar with NHC and quantified the amount of viral RNA released into the cell culture supernatant (Fig. 2B). Strikingly, the combination treatment diminished SARS-CoV-2 RNA progeny up to 400-fold as compared to single drug treatment, and up to 1000-fold as compared to untreated controls. This effect was not only seen in Vero E6 cells but also in Calu-3 cells (Fig. 2C), a human lung cancer cell line used to model bronchial epithelia (Kreft et al., 2015).

**FIGURE 2:**
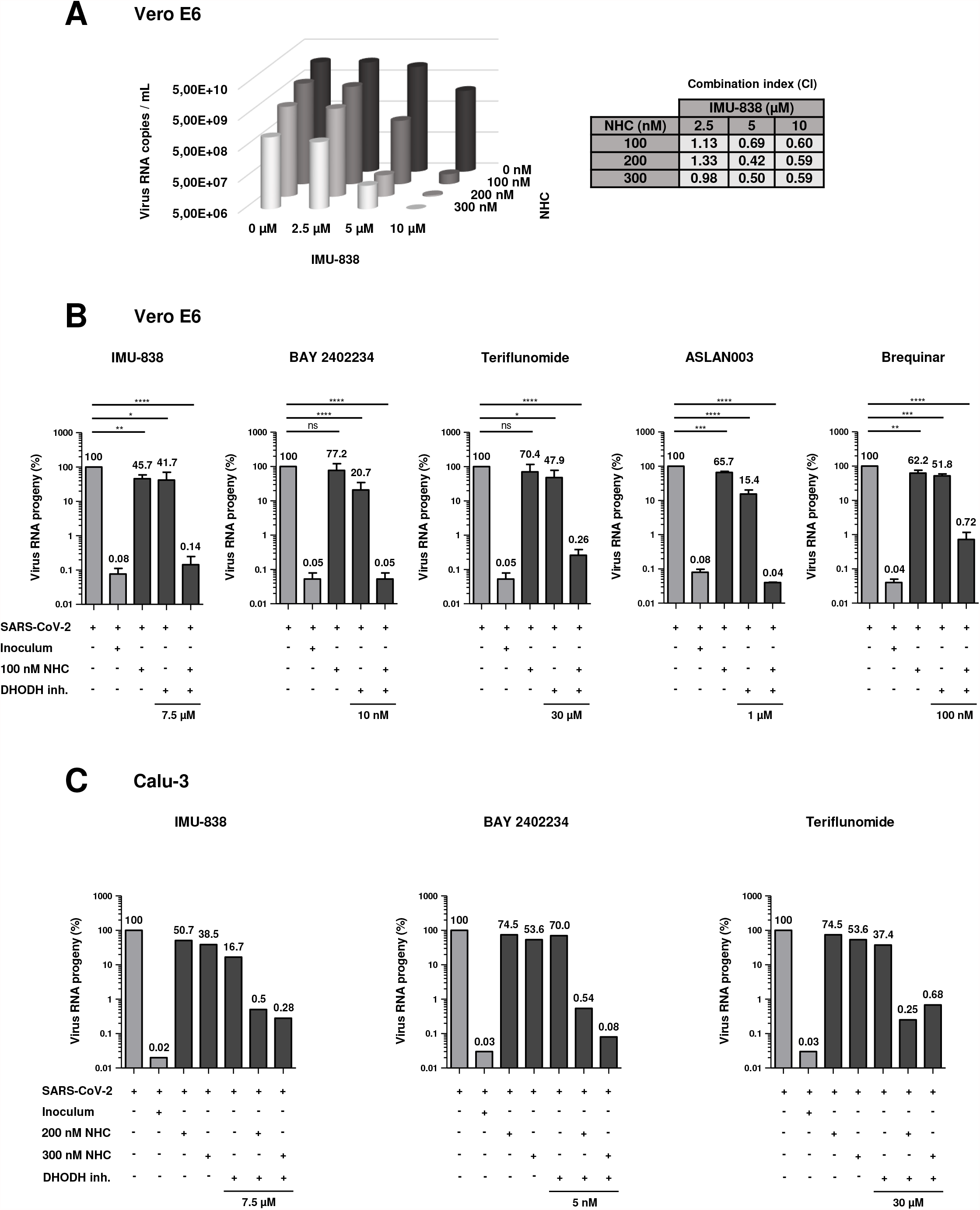
Strong synergism of NHC and DHODH inhibitors to diminish the release of SARS-CoV-2 RNA. **(A)** Reduced release of viral RNA by NHC and IMU-838. Vero E6 cells were treated with NHC and/or IMU-838 and/or infected as in Fig. 1. At 48 hrs post infection (p.i.), RNA was isolated from the cell supernatant, followed by quantitative RT-PCR to detect viral RNA and calculate the amount of SARS-CoV-2 RNA copies per mL (mean, n=3). The combination index (CI) was calculated using the CompuSyn software to determine the degree of drug synergism. **(B)** Reduced virus RNA progeny in the presence of NHC and DHODH inhibitors. Vero E6 cells were treated with drugs and/or infected, followed by quantitative detection of SARS-CoV-2 RNA. The amount of RNA found upon infection without drug treatment was defined as 100%, and the other RNA quantities were normalized accordingly. RNA was also isolated from the virus inoculum used to infect the cells. The drug combination was found capable of reducing virus RNA yield by more than 100-fold as compared to single drug treatments (mean with SD, n=3). **(C)** In Calu-3 cells, the combination of NHC and the DHODH inhibitors IMU-838, BAY2402234 or teriflunomide strongly reduced the amount of viral RNA released to the supernatant.

### Distinctly reduced accumulation of viral RNA in SARS-CoV-2 infected cells upon treatment with NHC and DHODH inhibitors

Next, we detected viral proteins upon treatment of Vero E6 cells with NHC and/or BAY2402234, teriflunomide or IMU-838, and infection with SARS-CoV-2. The frequency at which we detected the viral spike and nucleoprotein in the cells was severely reduced upon the combined treatment, and much less so by the single treatments, as determined by immunofluorescence microscopy (Fig. 3A-C). Correspondingly, immunoblot analysis revealed that both viral proteins were strongly diminished by the drug combination (Fig. 3D, E), whereas the single drugs at the same concentrations were much less efficient.

**FIGURE 3:**
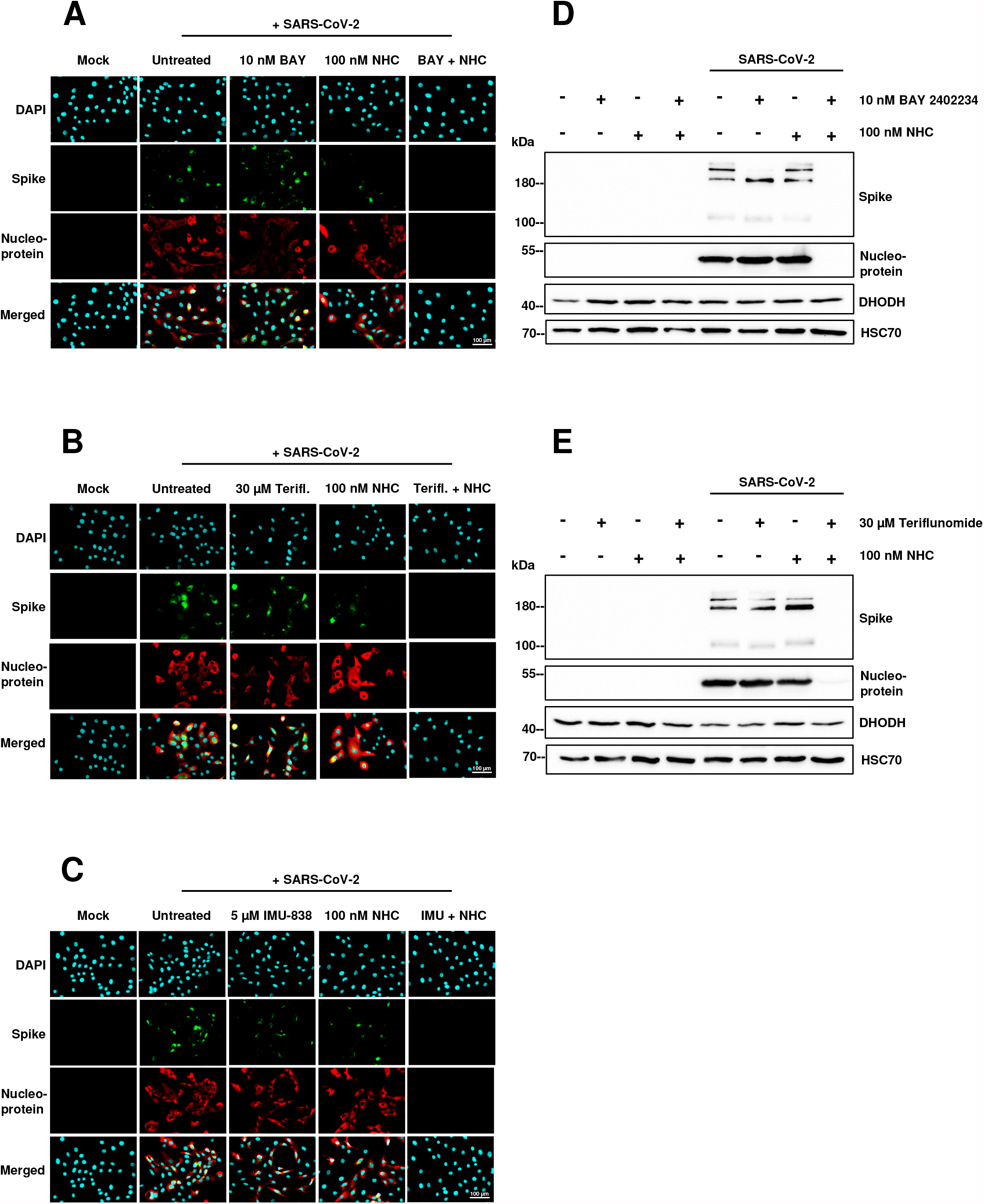
Synergistic reduction of viral protein synthesis by NHC and DHODH inhibitors. **(A, B and C)** Representative images showing the reduction of viral protein synthesis by NHC and BAY2402234 (A), teriflunomide (B), or IMU-838 (C). Vero E6 cells were treated as indicated and infected with SARS-CoV-2 as in Fig. 1. Cell nuclei were stained with DAPI, and the SARS-CoV-2 Spike and Nucleoprotein were detected by specific antibodies using immunofluorescence. **(D and E)** Reduced viral protein synthesis in the presence of NHC and DHODH inhibitors. Upon drug treatment and/or infection of Vero E6 cells as in Fig. 1, the viral Spike and Nucleoprotein as well as DHODH and HSC70 (loading control) were detected by immunoblot analysis.

### The combination of NHC and DHODH inhibitors also reduces the replication of the newly emerged SARS-CoV-2 variants B.1.1.7/Alpha and B.1.351/Beta

As the pandemic proceeded, new variants emerged, with potentially higher infectivity and immune-escape properties (Tegally et al., 2021; Wang et al., 2021). To ensure the suitability of the proposed drug combination against these variants, we assessed their replication in the presence of NHC and/or DHODH inhibitors. Both the variants B.1.1.7 (Alpha) and B.1.351 (Beta) responded similarly compared to the SARS-CoV-2 wildtype (Fig. 4A-C). Thus, the variations do not confer any detectable resistance against the drugs. This was expected since DHODH inhibitors target a cellular, not a viral function, i.e. pyrimidine synthesis, whereas even prolonged NHC treatment did not lead to resistance formation in other coronaviruses (Agostini et al., 2019).

**FIGURE 4:**
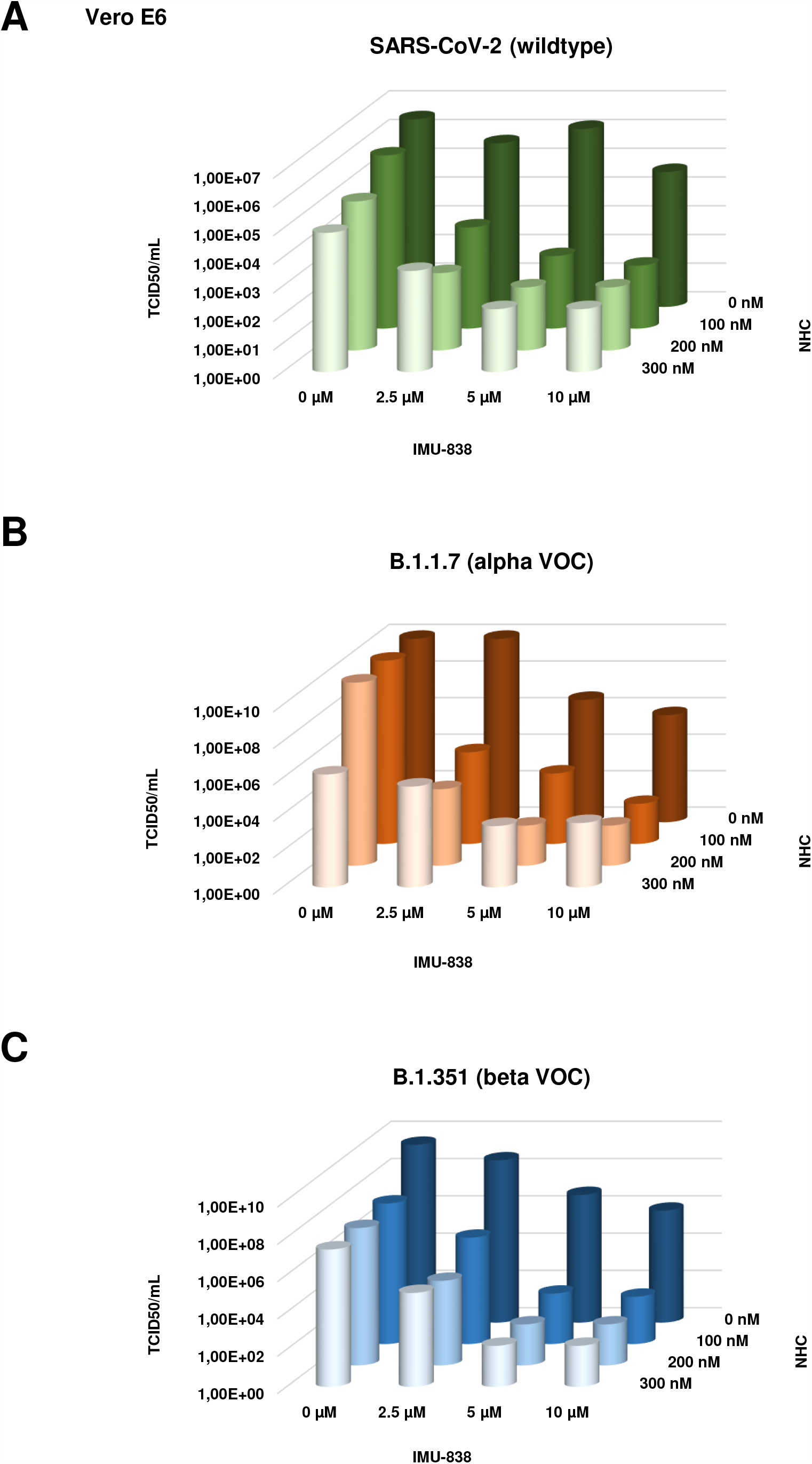
The combination of NHC with IMU-838 synergistically reduces the TCID_50_ of SARS-CoV-2 variants. **(A, B and C)** The TCID_50_ of virus progeny was determined upon treatment with NHC and DHODH inhibitors, upon infection with SARS-CoV-2 wildtype and two variants of concern (VOC). Vero E6 cells were treated with drugs for 24 hrs before and then throughout the infection. Cells were infected with SARS-CoV-2 (wildtype, (A)), B.1.1.7 (Alpha, (B)) or B.1.351 (Beta, (C)) (MOI 0.1) and further incubated for 48 hrs. The supernatant was titrated to determine the TCID_50_/mL. Note that both variants responded similarly to the wildtype, indicating that the drug combination is effective against newly emerged SARS-CoV-2 variants.

### Uridine and Cytidine rescue virus replication in the presence of NHC and DHODH inhibitors

To elucidate the mechanism of interference with SARS-CoV-2 replication by drug combinations, we performed rescue experiments by adding pyrimidine nucleosides. First, we added uridine to Vero E6 cells along with the DHODH inhibitors IMU-838, BAY2402234 or teriflunomide, combined with NHC (Fig. 5A). The addition of 2 µM uridine did not prevent the inhibition of SARS-CoV-2 replication by the combination treatment, but 10 µM uridine did. Similar results were obtained upon the addition of cytidine (Fig. 5B). This strongly suggests that the synergism of NHC with DHODH inhibitors can be explained by competition of NHC with endogenous pyrimidine nucleosides for incorporation into nascent viral RNA, as we had hypothesized initially (Fig. 1A).

**FIGURE 5:**
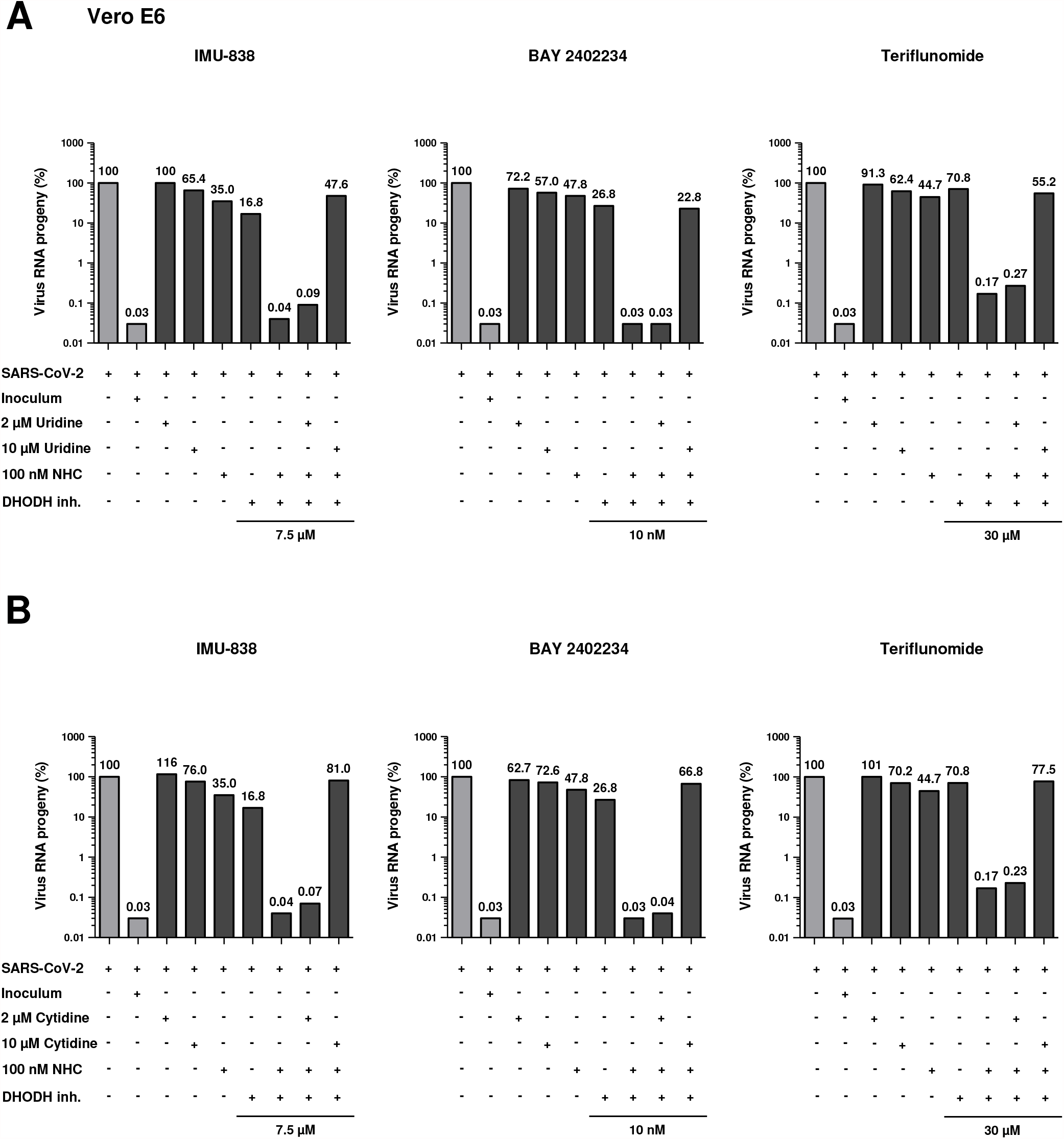
Uridine as well as cytidine rescue SARS-CoV-2 replication in the presence of NHC and DHODH inhibitors. **(A)** The antiviral effect of DHODH inhibitors combined with NHC can be rescued by uridine. Vero E6 cells were treated with drugs and inoculated with SARS-CoV-2 as shown in Fig. 1. On top of the drugs, where indicated, uridine was added to the cell culture media, at concentrations of 2 or 10 μM. SARS-CoV-2 propagation was still diminished by NHC and DHODH inhibitors despite 2 µM uridine levels, but rescued in the presence of 10 µM uridine. **(B)** Restored SARS-CoV-2 replication by cytidine, in the presence of NHC and DHODH inhibitors. The experiment was carried out as in (A), with the addition of cytidine instead of uridine. 10 μM cytidine restored virus replication in the presence of the drugs.

## DISCUSSION

Our results demonstrate that the simultaneous application of NHC and DHODH inhibitors suppresses the replication of SARS-CoV-2 far more profoundly than treatment with single drugs. Since both classes of compounds are undergoing advanced clinical evaluation for the treatment of COVID-19, our observations at least raise the perspective of using both drugs as antiviral combination therapy.

On top of enhancing the incorporation of NHC into viral RNA, DHODH inhibitors also suppress the immune response, e.g. by dampening the proliferation of B and T lymphocytes. In fact, they are currently in clinical use to treat autoimmune diseases (Munier-Lehmann et al., 2013). When treating COVID-19, a certain degree of immunosuppression may also be beneficial to avoid a “cytokine storm” (Fajgenbaum and June, 2020), as exemplified by the successful treatment with the steroid dexamethasone (Horby et al., 2021; Tomazini et al., 2020). When used alone, this bears the danger of re-enabling virus replication. However, we propose that the combination of DHODH inhibitors with NHC avoids both virus replication and hyperactive cytokine production.

Mechanistically, it is conceivable that reduced intracellular levels of cytidine triphosphate reduce the competition for triphosphorylated NHC regarding their incorporation into nascent virus RNA. This was further corroborated by the rescue of virus replication by uridine and cytidine, each metabolic precursors of CTP. Notably, the concentrations of uridine required for this rescue were in the range of 10 µM, which is above physiological serum concentrations (Ashour et al., 2000; Deng et al., 2017; Karle et al., 1980; Traut, 1994), raising the hope that pyrimidines in body fluids would not allow such rescue. NHC triphosphate is generated by the salvage pathway for pyrimidines but not by de novo pyrimidine synthesis, suggesting that NHC triphosphorylation is not impaired by DHODH inhibition. Thus, upon combined treatment, virus RNA will contain a larger proportion of NHC vs cytidine. In subsequent rounds of virus RNA replication, this will lead to misincorporations of adenine bases instead of guanine (Janion, 1978; Janion and Glickman, 1980; Salganik et al., 1973) with higher frequency, and render the virus genome nonfunctional due to missense and nonsense mutations.

In our previous work (Stegmann et al., 2021), we have established Methotrexate, a suppressor of purine biosynthesis, as an antagonist to SARS-CoV-2 replication, and this was confirmed and expanded recently (Caruso et al., 2021; Zhang et al., 2021). The principle behind this approach is similar to that of DHODH inhibitors, which also interfere with nucleotide biosynthesis. In our previous work, we also reported that Methotrexate cooperates with the antiviral purine analogue Remdesivir (Stegmann et al., 2021). However, this was not as pronounced as the synergy between NHC and DHODH inhibitors reported here. We speculate that even a relatively subtle increase in the ratio of NHC and cytidine in the template RNA strongly enhances the proportion of defective virions (Graci and Cameron, 2004, 2008), whereas the inhibitory effect of Remdesivir is only mildly enhanced by a reduction in available ATP. We also consider the possibility that the combined drugs might trigger signaling pathways that further interfere with virus replication. For instance, DHODH inhibition was found to affect the signaling mediators beta-catenin, p53 and MYC (Dorasamy et al., 2017; Qian et al., 2020), which might all affect the ability of a cell to support virus replication.

It remains to be determined how broadly this concept is applicable, i.e. combining inhibitors of pyrimidine synthesis with antiviral pyrimidine analogues. It is at least conceivable that such combinations can hinder the replication of other viruses as well, i.e. RNA and even DNA viruses. In fact, Molnupiravir was initially developed to counteract influenza virus replication (Toots et al., 2019; Toots et al., 2020). Even earlier, NHC was found to antagonize the propagation of Venezuelan equine encephalitis virus (VEEV) (Urakova et al., 2018) and coronavirus NL63 (Pyrc et al., 2006). Additional reports describe the impact of NHC on the replication of bovine viral diarrhea and hepatitis C virus (Stuyver et al., 2003), norovirus (Costantini et al., 2012), Ebola virus (Reynard et al., 2015), Chikungunya virus (Ehteshami et al., 2017), respiratory syncytial virus (Yoon et al., 2018), and the first pandemic sarbecovirus, i.e. SARS-CoV (Barnard et al., 2004). Likewise, DHODH inhibitors were found active against influenza virus as well, at least in vitro (Zhang et al., 2012). Other viruses susceptible to DHODH inhibition were negative-sense RNA viruses (besides influenza viruses A and B; Newcastle disease virus, and vesicular stomatitis virus), positive-sense RNA viruses (Sindbis virus, hepatitis C virus, West Nile virus, and dengue virus), DNA viruses (vaccinia virus and human adenovirus), and retroviruses (human immunodeficiency virus) (Hoffmann et al., 2011). Taken together, all these findings suggest that the combination of NHC and DHODH inhibitors might be successfully applicable to a broad range of viruses. This spectrum might be even further enhanced by combining inhibitors of pyrimidine synthesis with other pyrimidin-based antivirals, e.g. Sofosbuvir in treating hepatitis C virus infection (Manns et al., 2017), Lamivudine and Telbivudine against hepatitis B virus (Yuen et al., 2018), Brivudine to counteract varizella zoster virus (De Clercq, 2004), as well as several pyrimidine-based drugs against human immunodeficiency virus (Deeks et al., 2015).

NHC is a mutagen to bacteria (Janion, 1978; Janion and Glickman, 1980; Jena, 2020; Negishi et al., 1983; Popowska and Janion, 1974; Salganik et al., 1973). Presumably, NHC is converted to its 2’-deoxy form and then incorporated into bacterial DNA, followed by false-incorporation of adenine bases in the following round of DNA replication. It is currently unclear to what extent NHC induces mutagenesis in mammalian cells and possibly in patients when treated with its prodrug Molnupiravir. Mutagenesis of DNA is most likely determined by the 2’ reduction of NHC (in its diphosphorylated form) through deoxyribonucleotide reductase, by the degree of incorporation of deoxyNHC triphosphate through DNA polymerases, and by the efficiency of mismatch repair enzymes to replace NHC-induced mismatches with the correct C-G basepair. Among those, the ribonucleotide reductase activity would be druggable by currently available compounds, if necessary. Of note, however, Molnupiravir has not been reported to cause unacceptable levels of toxicities so far. Perhaps even more remarkably, ribavirin was not found cancerogenic after being used for decades in hepatitis C treatment, although its mechanism of action is also based on mutagenesis of virus RNA (Crotty et al., 2000). Thus, there is reason for cautious use of NHC-based drugs in the clinics, probably prohibiting their use during pregnancy; but in the face of ongoing pandemics, we still propose that mutagenesis in bacteria should not preclude the clinical evaluation of NHC and its prodrugs at least for the treatment of severe cases of COVID-19.

Remarkably, NHC treatment of coronavirus-infected cells did not give rise to resistant viruses, even after prolonged and repeated incubation (Agostini et al., 2019). Likewise, DHODH-inhibitors have a cellular rather than a viral target, thus providing little if any opportunity for resistant virus mutants to arise. Along with our finding that current virus variants of concern (VOC) respond similarly to the original SARS-CoV-2 strain (Fig. 4), this raises the hope that the drug combination will be universally applicable to treat most if not all SARS-CoV-2 variants, reducing disease burden and virus spread.

## ACKNOWLEDGEMENTS

We thank Tim Karrasch for help in isolating SARS-CoV-2-derived RNA and PCR-quantification. We thank Thorsten Wolff, Jessica Schulz and Christian Mache from the Robert-Koch Institute (Fachbereich 17) and Martin Beer from the Friedrich Loeffler Institute for providing Alpha (B.1.1.7) and Beta (B.1.351) isolates. Moreover, we thank Daniel Vitt and Hella Kohlhof, Immunic Therapeutics, for providing IMU-838 and for helpful advice, as well as Anne Balkema-Buschmann, Friedrich-Loeffler-Institute, for advising and critically reading the manuscript. KMS was a member of the Göttingen Graduate School GGNB during this work.

## AUTHOR CONTRIBUTIONS

MD conceived the project. KMS, AD, NH, and SP designed experiments. AD, KMS and NH performed experiments. SP, DG and UG provided guidance in virus isolation and quantification. MD and KMS wrote the manuscript. All authors revised and approved the manuscript.

## REFERENCES

Agostini, M.L., Pruijssers, A.J., Chappell, J.D., Gribble, J., Lu, X., Andres, E.L., Bluemling, G.R., Lockwood, M.A., Sheahan, T.P., Sims, A.C., et al. (2019). Small-Molecule Antiviral β-d-N (4)-Hydroxycytidine Inhibits a Proofreading-Intact Coronavirus with a High Genetic Barrier to Resistance. Journal of virology 93.

Ashour, O.M., Al Safarjalani, O.N., Naguib, F.N., Goudgaon, N.M., Schinazi, R.F., and el Kouni, M.H. (2000). Modulation of plasma uridine concentration by 5-(phenylselenenyl)acyclouridine, an inhibitor of uridine phosphorylase: relevance to chemotherapy. Cancer chemotherapy and pharmacology 45, 351–361.

Barnard, D.L., Hubbard, V.D., Burton, J., Smee, D.F., Morrey, J.D., Otto, M.J., and Sidwell, R.W. (2004). Inhibition of severe acute respiratory syndrome-associated coronavirus (SARSCoV) by calpain inhibitors and beta-D-N4-hydroxycytidine. Antiviral chemistry & chemotherapy 15, 15–22.

Beigel, J.H., Tomashek, K.M., Dodd, L.E., Mehta, A.K., Zingman, B.S., Kalil, A.C., Hohmann, E., Chu, H.Y., Luetkemeyer, A., Kline, S., et al. (2020). Remdesivir for the Treatment of Covid-19 - Final Report. The New England journal of medicine 383, 1813–1826.

Caruso, A., Caccuri, F., Bugatti, A., Zani, A., Vanoni, M., Bonfanti, P., Cazzaniga, M.E., Perno, C.F., Messa, C., and Alberghina, L. (2021). Methotrexate inhibits SARS-CoV-2 virus replication “in vitro”. Journal of medical virology 93, 1780–1785.

Chou, T.C. (2010). Drug combination studies and their synergy quantification using the Chou-Talalay method. Cancer research 70, 440–446.

Corman, V.M., Landt, O., Kaiser, M., Molenkamp, R., Meijer, A., Chu, D.K., Bleicker, T., Brünink, S., Schneider, J., Schmidt, M.L., et al. (2020). Detection of 2019 novel coronavirus (2019-nCoV) by real-time RT-PCR. Euro surveillance : bulletin Europeen sur les maladies transmissibles = European communicable disease bulletin 25.

Costantini, V.P., Whitaker, T., Barclay, L., Lee, D., McBrayer, T.R., Schinazi, R.F., and Vinjé, J. (2012). Antiviral activity of nucleoside analogues against norovirus. Antiviral therapy 17, 981–991.

Cox, R.M., Wolf, J.D., and Plemper, R.K. (2021). Therapeutically administered ribonucleoside analogue MK-4482/EIDD-2801 blocks SARS-CoV-2 transmission in ferrets. Nat Microbiol 6, 11–18.

Crotty, S., Maag, D., Arnold, J.J., Zhong, W., Lau, J.Y.N., Hong, Z., Andino, R., and Cameron, C.E. (2000). The broad-spectrum antiviral ribonucleoside ribavirin is an RNA virus mutagen. Nature Medicine 6, 1375–1379.

De Clercq, E. (2004). Discovery and development of BVDU (brivudin) as a therapeutic for the treatment of herpes zoster. Biochemical pharmacology 68, 2301–2315.

Deeks, S.G., Overbaugh, J., Phillips, A., and Buchbinder, S. (2015). HIV infection. Nature Reviews Disease Primers 1, 15035.

Deng, Y., Wang, Z.V., Gordillo, R., An, Y., Zhang, C., Liang, Q., Yoshino, J., Cautivo, K.M., De Brabander, J., Elmquist, J.K., et al. (2017). An adipo-biliary-uridine axis that regulates energy homeostasis. Science (New York, N.Y.) 355, eaaf5375.

Doherty, P.C. (2021). What have we learnt so far from COVID-19? Nature Reviews Immunology 21, 67–68.

Dorasamy, M.S., Choudhary, B., Nellore, K., Subramanya, H., and Wong, P.F. (2017). Dihydroorotate dehydrogenase Inhibitors Target c-Myc and Arrest Melanoma, Myeloma and Lymphoma cells at S-phase. Journal of Cancer 8, 3086–3098.

Ehteshami, M., Tao, S., Zandi, K., Hsiao, H.M., Jiang, Y., Hammond, E., Amblard, F., Russell, O.O., Merits, A., and Schinazi, R.F. (2017). Characterization of β-d-N(4)-Hydroxycytidine as a Novel Inhibitor of Chikungunya Virus. Antimicrobial agents and chemotherapy 61.

Fajgenbaum, D.C., and June, C.H. (2020). Cytokine Storm. The New England journal of medicine 383, 2255–2273.

Goldman, J.D., Lye, D.C.B., Hui, D.S., Marks, K.M., Bruno, R., Montejano, R., Spinner, C.D., Galli, M., Ahn, M.Y., Nahass, R.G., et al. (2020). Remdesivir for 5 or 10 Days in Patients with Severe Covid-19. The New England journal of medicine 383, 1827–1837.

Graci, J.D., and Cameron, C.E. (2004). Challenges for the development of ribonucleoside analogues as inducers of error catastrophe. Antiviral chemistry & chemotherapy 15, 1–13.

Graci, J.D., and Cameron, C.E. (2008). Therapeutically targeting RNA viruses via lethal mutagenesis. Future virology 3, 553–566.

Hahn, F., Wangen, C., Hage, S., Peter, A.S., Dobler, G., Hurst, B., Julander, J., Fuchs, J., Ruzsics, Z., Uberla, K., et al. (2020). IMU-838, a Developmental DHODH Inhibitor in Phase II for Autoimmune Disease, Shows Anti-SARS-CoV-2 and Broad-Spectrum Antiviral Efficacy In Vitro. Viruses 12.

Hoffmann, H.H., Kunz, A., Simon, V.A., Palese, P., and Shaw, M.L. (2011). Broad-spectrum antiviral that interferes with de novo pyrimidine biosynthesis. Proceedings of the National Academy of Sciences of the United States of America 108, 5777–5782.

Horby, P., Lim, W.S., Emberson, J.R., Mafham, M., Bell, J.L., Linsell, L., Staplin, N., Brightling, C., Ustianowski, A., Elmahi, E., et al. (2021). Dexamethasone in Hospitalized Patients with Covid-19. The New England journal of medicine 384, 693–704.

Janion, C. (1978). The efficiency and extent of mutagenic activity of some new mutagens of base-analogue type. Mutation research 56, 225–234.

Janion, C., and Glickman, B.W. (1980). N4-hydroxycytidine: a mutagen specific for AT to GC transitions. Mutation research 72, 43–47.

Jena, N.R. (2020). Role of different tautomers in the base-pairing abilities of some of the vital antiviral drugs used against COVID-19. Physical chemistry chemical physics : PCCP 22, 28115–28122.

Karle, J.M., Anderson, L.W., Erlichman, C., and Cysyk, R.L. (1980). Serum uridine levels in patients receiving N-(phosphonacetyl)-L-aspartate. Cancer research 40, 2938–2940.

Kreft, M.E., Jerman, U.D., Lasič, E., Hevir-Kene, N., Rižner, T.L., Peternel, L., and Kristan, K. (2015). The characterization of the human cell line Calu-3 under different culture conditions and its use as an optimized in vitro model to investigate bronchial epithelial function. European journal of pharmaceutical sciences : official journal of the European Federation for Pharmaceutical Sciences 69, 1–9.

Luban, J., Sattler, R.A., Mühlberger, E., Graci, J.D., Cao, L., Weetall, M., Trotta, C., Colacino, J.M., Bavari, S., Strambio-De-Castillia, C., et al. (2021). The DHODH inhibitor PTC299 arrests SARS-CoV-2 replication and suppresses induction of inflammatory cytokines. Virus Res 292, 198246.

Manns, M.P., Buti, M., Gane, E., Pawlotsky, J.-M., Razavi, H., Terrault, N., and Younossi, Z. (2017). Hepatitis C virus infection. Nature Reviews Disease Primers 3, 17006.

Munier-Lehmann, H., Vidalain, P.O., Tangy, F., and Janin, Y.L. (2013). On dihydroorotate dehydrogenases and their inhibitors and uses. Journal of medicinal chemistry 56, 3148–3167.

Negishi, K., Harada, C., Ohara, Y., Oohara, K., Nitta, N., and Hayatsu, H. (1983). N4-aminocytidine, a nucleoside analog that has an exceptionally high mutagenic activity. Nucleic acids research 11, 5223–5233.

Painter, W.P., Holman, W., Bush, J.A., Almazedi, F., Malik, H., Eraut, N., Morin, M.J., Szewczyk, L.J., and Painter, G.R. (2021). Human Safety, Tolerability, and Pharmacokinetics of Molnupiravir, a Novel Broad-Spectrum Oral Antiviral Agent with Activity Against SARS-CoV-2. Antimicrobial agents and chemotherapy.

Popowska, E., and Janion, C. (1974). N4-hydroxycytidine-a new mutagen of a base analogue type. Biochemical and biophysical research communications 56, 459–466.

Pruijssers, A.J., and Denison, M.R. (2019). Nucleoside analogues for the treatment of coronavirus infections. Current opinion in virology 35, 57–62.

Pyrc, K., Bosch, B.J., Berkhout, B., Jebbink, M.F., Dijkman, R., Rottier, P., and van der Hoek, L. (2006). Inhibition of human coronavirus NL63 infection at early stages of the replication cycle. Antimicrobial agents and chemotherapy 50, 2000–2008.

Qian, Y., Liang, X., Kong, P., Cheng, Y., Cui, H., Yan, T., Wang, J., Zhang, L., Liu, Y., Guo, S., et al. (2020). Elevated DHODH expression promotes cell proliferation via stabilizing β-catenin in esophageal squamous cell carcinoma. Cell Death & Disease 11, 862.

Reynard, O., Nguyen, X.N., Alazard-Dany, N., Barateau, V., Cimarelli, A., and Volchkov, V.E. (2015). Identification of a New Ribonucleoside Inhibitor of Ebola Virus Replication. Viruses 7, 6233–6240.

Salganik, R.I., Vasjunina, E.A., Poslovina, A.S., and Andreeva, I.S. (1973). Mutagenic action of N4-hydroxycytidine on Escherichia coli B cyt. Mutation research 20, 1–5.

Sheahan, T.P., Sims, A.C., Zhou, S., Graham, R.L., Pruijssers, A.J., Agostini, M.L., Leist, S.R., Schafer, A., Dinnon, K.H., 3rd, Stevens, L.J., et al. (2020). An orally bioavailable broad-spectrum antiviral inhibits SARS-CoV-2 in human airway epithelial cell cultures and multiple coronaviruses in mice. Science translational medicine 12.

Stegmann, K.M., Dickmanns, A., Gerber, S., Nikolova, V., Klemke, L., Manzini, V., Schlösser, D., Bierwirth, C., Freund, J., Sitte, M., et al. (2021). The folate antagonist methotrexate diminishes replication of the coronavirus SARS-CoV-2 and enhances the antiviral efficacy of remdesivir in cell culture models. Virus research 302, 198469.

Stuyver, L.J., Whitaker, T., McBrayer, T.R., Hernandez-Santiago, B.I., Lostia, S., Tharnish, P.M., Ramesh, M., Chu, C.K., Jordan, R., Shi, J., et al. (2003). Ribonucleoside analogue that blocks replication of bovine viral diarrhea and hepatitis C viruses in culture. Antimicrobial agents and chemotherapy 47, 244–254.

Tegally, H., Wilkinson, E., Giovanetti, M., Iranzadeh, A., Fonseca, V., Giandhari, J., Doolabh, D., Pillay, S., San, E.J., Msomi, N., et al. (2021). Detection of a SARS-CoV-2 variant of concern in South Africa. Nature 592, 438–443.

Tomazini, B.M., Maia, I.S., Cavalcanti, A.B., Berwanger, O., Rosa, R.G., Veiga, V.C., Avezum, A., Lopes, R.D., Bueno, F.R., Silva, M., et al. (2020). Effect of Dexamethasone on Days Alive and Ventilator-Free in Patients With Moderate or Severe Acute Respiratory Distress Syndrome and COVID-19: The CoDEX Randomized Clinical Trial. Jama 324, 1307–1316.

Toots, M., Yoon, J.J., Cox, R.M., Hart, M., Sticher, Z.M., Makhsous, N., Plesker, R., Barrena, A.H., Reddy, P.G., Mitchell, D.G., et al. (2019). Characterization of orally efficacious influenza drug with high resistance barrier in ferrets and human airway epithelia. Science translational medicine 11.

Toots, M., Yoon, J.J., Hart, M., Natchus, M.G., Painter, G.R., and Plemper, R.K. (2020). Quantitative efficacy paradigms of the influenza clinical drug candidate EIDD-2801 in the ferret model. Translational research : the journal of laboratory and clinical medicine 218, 16–28.

Traut, T.W. (1994). Physiological concentrations of purines and pyrimidines. Molecular and Cellular Biochemistry 140, 1–22.

Urakova, N., Kuznetsova, V., Crossman, D.K., Sokratian, A., Guthrie, D.B., Kolykhalov, A.A., Lockwood, M.A., Natchus, M.G., Crowley, M.R., Painter, G.R., et al. (2018). beta-d-N (4)-Hydroxycytidine Is a Potent Anti-alphavirus Compound That Induces a High Level of Mutations in the Viral Genome. Journal of virology 92.

Wahl, A., Gralinski, L., Johnson, C., Yao, W., Kovarova, M., Dinnon, K., Liu, H., Madden, V., Krzystek, H., De, C., et al. (2020). Acute SARS-CoV-2 Infection is Highly Cytopathic, Elicits a Robust Innate Immune Response and is Efficiently Prevented by EIDD-2801. Res Sq.

Wahl, A., Gralinski, L.E., Johnson, C.E., Yao, W., Kovarova, M., Dinnon, K.H., 3rd, Liu, H., Madden, V.J., Krzystek, H.M., De, C., et al. (2021). SARS-CoV-2 infection is effectively treated and prevented by EIDD-2801. Nature 591, 451–457.

Wang, P., Nair, M.S., Liu, L., Iketani, S., Luo, Y., Guo, Y., Wang, M., Yu, J., Zhang, B., Kwong, P.D., et al. (2021). Antibody resistance of SARS-CoV-2 variants B.1.351 and B.1.1.7. Nature.

Xiong, R., Zhang, L., Li, S., Sun, Y., Ding, M., Wang, Y., Zhao, Y., Wu, Y., Shang, W., Jiang, X., et al. (2020). Novel and potent inhibitors targeting DHODH are broad-spectrum antivirals against RNA viruses including newly-emerged coronavirus SARS-CoV-2. Protein Cell 11, 723–739.

Yoon, J.J., Toots, M., Lee, S., Lee, M.E., Ludeke, B., Luczo, J.M., Ganti, K., Cox, R.M., Sticher, Z.M., Edpuganti, V., et al. (2018). Orally Efficacious Broad-Spectrum Ribonucleoside Analog Inhibitor of Influenza and Respiratory Syncytial Viruses. Antimicrobial agents and chemotherapy 62.

Yuen, M.-F., Chen, D.-S., Dusheiko, G.M., Janssen, H.L.A., Lau, D.T.Y., Locarnini, S.A., Peters, M.G., and Lai, C.-L. (2018). Hepatitis B virus infection. Nature Reviews Disease Primers 4, 18035.

Zhang, L., Das, P., Schmolke, M., Manicassamy, B., Wang, Y., Deng, X., Cai, L., Tu, B.P., Forst, C.V., Roth, M.G., et al. (2012). Inhibition of pyrimidine synthesis reverses viral virulence factor-mediated block of mRNA nuclear export. The Journal of cell biology 196, 315–326.

Zhang, Y., Guo, R., Kim, S.H., Shah, H., Zhang, S., Liang, J.H., Fang, Y., Gentili, M., Leary, C.N.O., Elledge, S.J., et al. (2021). SARS-CoV-2 hijacks folate and one-carbon metabolism for viral replication. Nature Communications 12, 1676.

